# Interactions between the gut microbiome and host gene regulation in cystic fibrosis

**DOI:** 10.1101/596312

**Authors:** Gargi Dayama, Sambhawa Priya, David E. Niccum, Alexander Khoruts, Ran Blekhman

**Author notes:** these authors contributed equally (Gargi Dayama, Sambhawa Priya). to whom correspondence should be addressed (AK), (RB).

## Abstract

Cystic Fibrosis (CF) is the most common autosomal recessive genetic disease in Caucasians. It is caused by mutations in the *CFTR* gene, leading to poor hydration of mucus and impairment of the respiratory, digestive, and reproductive organ functions. Advancements in medical care have lead to markedly increased longevity of patients with CF, but new complications have emerged, such as early onset of colorectal cancer (CRC). Although the pathogenesis of CRC in CF remains unclear, altered host-microbe interactions might play a critical role. Here, we characterize the changes in the gut microbiome and host gene expression in colonic mucosa of CF patients relative to healthy controls. We find that CF patients show decreased microbial diversity, decreased abundance of taxa such as *Butyricimonas, Sutterella,* and Ruminococcaceae, and increased abundance of other taxa, such as Actinobacteria and Firmicutes. We find that 1543 genes, including *CFTR,* show differential expression in the colon of CF patients compared to healthy controls. Interestingly, we find that these genes are enriched with functions related to gastrointestinal and colorectal cancer, such as metastasis of CRC, tumor suppression, cellular dysfunction, p53 and mTOR signaling pathways. Lastly, we modeled associations between relative abundances of specific bacterial taxa in the gut mucosa and host gene expression, and identified CRC-related genes, including *LCN2* and *DUOX2,* for which gene expression is correlated with the abundance of CRC-associated bacteria, such as Ruminococcaceae and *Veillonella*. Our results provide new insight into the role of host-microbe interactions in the etiology of CRC in CF.

## Introduction

Cystic fibrosis (CF) is the most common autosomal recessive genetic disease in Caucasians, where it occurs with a frequency of 1 in 3,000 births, although it is also present at lower rates in populations of non-European descent [1]. CF is caused by mutations in the cystic fibrosis transmembrane conductor regulatory *(CFTR)* gene, which plays critical functions in epithelial ion transport and hydration of mucus. Absent or reduced *CFTR* activity results in thick, viscous secretions that impair functions of the respiratory, digestive, and reproductive organ systems.

Multiple advances in medical care in CF, once a fatal pediatric disease, have led to remarkable gains in patient life expectancy. However, increased longevity of CF patients into adulthood has led to new challenges, such as gastrointestinal cancer. The average onset of colorectal cancer (CRC) in CF patients is approximately 20-30 years earlier than in the general population [2,3]. Systematic data on colonoscopic screening and surveillance suggest that CF-associated CRC arises via the classical adenoma to cancer sequence, but adenomatous polyps develop at a younger age in CF and progress faster to more advanced neoplasms [4]. In fact, loss of *CFTR* expression in tumors of non-CF patients has been associated with a worse prognosis in early stage CRC [5]. Recently, specific recommendations for CRC screening were introduced in standard care of adult CF patients, which include earlier initiation of screening and shorter intervals for surveillance [6].

Although previous studies have identified *CFTR* as a tumor suppressor gene that may play a role in early onset of colon cancer [5,7], the pathogenesis of CRC in CF remains unclear. A number of factors can be considered. Thus, stagnant mucus in CF is associated with bacterial overgrowth at the mucosal surface [8,9], which might result in greater levels of tonic microbial stimulation of the epithelia and account for their increased rate of their turnover [10]. It is likely that the altered microbiota composition and microbiota-mucosal interface are also the reasons for a chronic state of low-grade mucosal inflammation in CF [11,12]. Notably, in the colon *CFTR* is hyper-expressed in the stem cell compartment of the intestinal crypt [13,14], which is the site of CRC origination [15].

Than and colleagues have shown altered expression of genes involved in immune cell homeostasis and inflammation, mucins, cell signaling and growth regulation, detoxification and stress response, lipid metabolism, and stem cell regulation in the intestines of *CFTR* mutant mice [5]. The intestinal microbiota of these animals is also distinguished by lower bacterial community richness, evenness, and diversity, consistent with a major impact of *CFTR* deficiency on gastrointestinal physiology [16]. Altered fecal microbiome has also been demonstrated in a number of clinical CF cohorts, where it was characterized by decreased microbial diversity, lower temporal microbial community stability, and decreased relative abundances of taxa associated with health, such as *Faecalibacterium, Roseburia, Bifidobacterium, Akkermansia, Clostridium cluster XIVa* [17–23]. Greater degrees of dysbiosis were noted to correlate with severity of CF disease phenotype, burden of antibiotics, and evidence for intestinal inflammation. Notably, most of these patient studies have been in diverse pediatric cohorts with varying degrees of fat malabsorption and extent of recent exposure to broad-spectrum antibiotics.

Here, we compare the mucosal microbiome (via 16S rRNA sequencing) and colonic gene expression (via RNA-seq) in adult patients with CF and healthy controls undergoing CRC screening by colonoscopy. We explore interactions between the gut microbiome and host gene regulation by integrative analysis, characterizing genes and microbes that may play a joint role in the development of CRC in CF patients.

## Methods

### Patients and mucosal biopsy samples

Mucosal biopsies were obtained from patients undergoing CRC screening and surveillance colonoscopies at the University of Minnesota. The majority of CF patients receiving care at the Minnesota Cystic Fibrosis Center participate in a systematic colonoscopic CRC screening program as described previously [4]. Control samples were obtained from non-CF patients undergoing routine colonoscopic CRC screening or surveillance. Pinch biopsies, four per patient, were obtained using the Radial Jaw 4 Jumbo w/Needle 240 (length) forceps for 3.2 mm working channel (Boston Scientific, Marlborough, MA; Catalog # M00513371) in the right colon and placed into RNAlater stabilization solution (ThermoFisher Scientific, Waltham, MA). The protocol was approved by the University of Minnesota Institutional Review Board (IRB protocol 1408M52889). Gene expression was analyzed by RNA-Seq from a total of 33 samples obtained from 18 CF patients and 15 non-CF control participants (Fig S1).

### RNA extraction and sequencing

Biopsy tissue was kept in the RNAlater stabilization solution overnight at 4°C. RNA was prepared following tissue homogenization and lysis using the TRIzol Plus RNA Purification Kit (ThermoFisher Scientific; catalogue # 2183-555) following detailed manufacturer instructions. Total RNA samples were converted to Illumina sequencing libraries using Illumina’s Truseq Stranded mRNA Sample Preparation Kit (Cat. # RS-122-2103). Total RNA is oligo-dT purified using oligo-dT coated magnetic beads, fragmented and then reverse transcribed into cDNA. The cDNA is adenylated and then ligated to dual-indexed (barcoded) adaptors and amplified using 15 cycles of PCR. Final library size distribution is validated using capillary electrophoresis and quantified using fluorimetry (PicoGreen). Indexed libraries are then normalized, pooled and then size selected to 320bp +/− 5% using Caliper’s XT instrument. Truseq libraries are hybridized to a paired end flow cell and individual fragments are clonally amplified by bridge amplification on the Illumina cBot. Once clustering is complete, the flow cell is loaded on the HiSeq 2500 and sequenced using Illumina’s SBS chemistry (Fig S1).

### Host RNA-seq quality control, read mapping and filtering

We performed quality check on raw sequences from all 33 samples to assure better downstream analysis using FastQC [24]. This helped assess any biases due to parameters such as quality of the reads, GC content, number of reads, read length and species to which the majority of the reads mapped (Fig S2). The FASTQ files for forward and reverse (R1 and R2) reads were mapped to the reference genome using kallisto [25], where an index for the transcriptomes was generated to quantify estimated read counts and TPM values. Mean distribution for the TPM values was plotted using R to filter all the transcripts below a threshold value of log2[TPM] < 0. We generated PCA plots using sleuth [26] to examine sample clusters and visualization of expression patterns for genes using bar plots (Fig S3 and Fig S4). For further analysis of outlier samples box plots were generated using Cook’s distance and heat map clustered by condition and mutation status was generated for the top 20 expressed genes (Fig S5 and Fig S6).

### Host RNA-seq differential expression and enrichment analysis

To determine differentially expressed genes between CF and healthy samples we quantified and annotated the transcripts used DESeq2 [27]. The output from kallisto was imported into DESeq2 using the tximport package [28]. The transcripts were annotated against the ensemble database using bioMART to obtain gene symbols [29].

Transcripts below a threshold of row-sum of 1 were filtered and collapsed at gene symbol level. Prior to differentially expressed gene analysis, the read counts were normalized and the gene-wise estimates were shrunken towards the fitted estimates represented by the red line in the dispersion plot (Fig S7). The gene-wise estimates that are outliers are not shrunk and are flagged by the blue circles in the plot (Fig S7). DESeq2 applies the Wald test on estimated counts and uses a negative binomial generalized linear model determines differentially expressed genes and the log-fold changes (Fig S8). The log-fold change shrinkage *(lcfshrink())* function was applied for ranking the genes and data visualization. For data smoothing, MA plots were generated before and after log2 fold shrinkage. We found no change in the MA plot (Fig S9) post smoothing, as there are no large log-fold changes in the current data (log2 fold change between −1 and 1) due to low counts. The data were further transformed and the normalized values were extracted using regularized logarithm (rlog) to remove the dependence of variance on mean. We used the Benjamini-Hochberg method for reducing the false discovery rate (FDR) with a cutoff of 0.05 for identifying differentially expressed genes for further analysis. Enrichment analysis was done using Ingenuity Pathway Analysis (IPA, QIAGEN Inc., https://www.qiagenbioinformatics.com/products/ingenuitypathway-analysis). The logfold changes, p-values, and FDR values (for all the genes with FDR < 0.05) were fed into IPA for both up- and down-regulated differentially expressed genes between CF and healthy samples. Disease/functional pathways and gene networks were determined based on the gene enrichment. Furthermore, we looked at how many target upstream regulators were enriched based on our list of differentially expressed genes using IPA. We found 134 targets that passed the filter (p-value < 0.01) from a total of 492 targets, of which 96 were transcription regulators.

### 16S rRNA extraction and sequencing

Mucosal biopsies samples (~ 3 × 3 mm) from 13 CF and 12 healthy individuals were collected in 1 mL of RNAlater and stored for 24 hours at 4°C prior to freezing at −80°C. DNA was extracted using a MoBio PowerSoil DNA isolation kit according to the manufacturer instructions (QIAGEN, Carlsbad, USA). To look at the tissue associated microbiome, V5-V6 region of 16S rRNA gene was amplified as described by Huse et al. [30] using the following indexing primers (V5F_Nextera: TCGTCGGCAGCGTCAGATGTGTATAAGAGACAGRGG ATTAGATACCC, V6R_Nextera: GTCTCGTGGGCTCGGAGATGTGTATAAGAGACAGCGACRRCCATGCANCACCT). Index and flowcell adaptors were added with this step. Forward indexing primer used is – **AATGATACGGCGACCACCGA**GATCTACAC[i5]TCGTCGGCAGCGTC and reverse indexing primers used is – **CAAGCAGAAGACGGCATACGA**GAT[i7]GTCTCGTGGGCTCGG. Post two rounds of PCR, pooled, size-selected samples were denatured with NaOH, diluted to 8 pM in Illumina’s HT1 buffer, spiked with 15% PhiX, and heat denatured at 96°C for 2 minutes immediately prior to loading. A MiSeq 600 cycle v3 kit was used to sequence the sample.

### Gut mucosal microbiome data processing, quality assessment, and diversity analysis

We processed the FASTQ files using FastQC [24] to perform quality control on the raw sequences. We then used SHI7 [31] for trimming Nextera adaptors, stitching paired-end reads and performing quality trimming at both ends of the stitched reads until a minimum Phred score of 32 was reached. Following quality control, we obtained an average of 217,500 high quality reads per sample (median 244,000; range 9551 – 373,900) with an average length of 281.9 bases and an average quality score of 37.19. These merged and filtered reads were used for closed reference OTU picking and taxonomy assignment against GreenGenes database with 97% similarity level using the NINJA-OPS program [32].

We performed alpha and beta-diversity analysis in R using the vegan [33] and phyloseq [34] packages. We used resampling-based computation of alpha-diversity, where the OTU table is subsampled 100 times at minimum read depth (9551 reads) across all samples, and computed average richness estimate for each alpha-diversity metric (chao1, observed-OTUs, and Shannon). Wilcoxon rank-sum test was used for testing the statistical significance of the associations between alpha-diversity of the CF and healthy conditions. For computing beta-diversity, we first rarefied the OTU table (using vegan’s *rrarefy()* function) at minimum sequence depth (i.e. 9551 reads) across the samples and then computed Bray-Curtis dissimilarity, weighted UniFrac, and unweighted UniFrac metrics. The Adonis test was used for assessing if there is significant association between the beta-diversity of the CF/healthy condition and the diversity results are plotted using the ggplot2 package in R.

### Gut mucosal microbiome differential abundance and functional analysis

We performed differential abundance testing using the phyloseq [34] package in R. We first created a phyloseq object from the OTU table (using the *phyloseq()* function) and filtered this object to only include OTUs with at least 0.1% relative abundance occurring in at least half of all samples (using the *filter_taxa()* function). The filtered phyloseq object was converted into a DESeqDataSet object (using *phyloseq_to_deseq2())* and the *DESeq()* function was invoked. This performed dispersion estimations and Wald’s test for identifying differentially abundant OTUs, with their corresponding log-fold change, p-value, and FDR-adjusted q-values between the CF and healthy conditions. We agglomerated the OTUs at different taxonomic ranks (using the *tax_glom()* function) and repeated the above steps to identify differentially abundant taxa at genus, family, order, class, and phylum levels.

We also tested for associations between taxonomic abundance and mutation status of CF samples. We first categorized samples into three genotype categories: (1) Healthy: Samples with no mutations; (2) CF_df508: CF samples with homozygous delta-F508 deletion, which is associated with more severe CF condition [35]; and (3) CF_other: CF samples with df508 heterozygous deletion or other mutation status. We used DESeq2’s likelihood ratio test (LRT) to identify taxa that showed significant difference in abundance across the three categories.

We then generated the predicted functional profiles for the gut microbes using PICRUSt v1.0.0 pipeline, [36] where pathways and enzymes are assigned using the Kyoto Encyclopedia of Genes and Genomes (KEGG) database. The KEGG level 3 pathways were filtered for rare pathways by only including pathways with relative abundance > 0.1% in at least half of the samples, normalized to relative abundance, and tested for association with CF/Healthy conditions using non-parametric Wilcoxon rank-sum test followed by FDR adjustment.

### Integrated analysis of interactions between host gene dysregulations and changes in microbiome

For this analysis, differentially expressed genes from host and gut microbial OTUs from their respective overlapping samples were used (22 samples in total, with 12 healthy samples and 10 CF samples). We further subset differentially expressed genes between CF and healthy conditions (FDR < 0.05), specifically, enriched for gastrointestinal cancer disease pathways (524 genes). Using absolute expression log ratio greater than 0.35, we obtained a representative set of both up- and down-regulated genes from these pathways, leaving 250 genes for downstream analysis. The OTU table was collapsed at the genus level (or the last characterized level) and filtered for rare taxa by only including taxa with at least 0.1% relative abundance present in at least half of all samples, resulting in 35 taxa for further processing. Following this, centered log ratio transform was applied on the filtered table. We then performed correlation analysis between host gene expression data for 250 genes and gut microbiome abundance data for 35 taxa (genus level) defined above. Spearman correlation was used for this analysis as it performs better with normalized counts (gene expression) as well as compositional data (microbiome relative abundance) compared to other metrics, such as Pearson correlation (Weiss et al. 2016). We computed the Spearman rank correlation coefficients and the corresponding p-values using the *cor.test()* function with two-sided alternative hypothesis. A total of 8750 (250 genes x 35 taxa) statistical tests were performed, and p-values were corrected for multiple comparisons using the qvalue package in R [37]. Representative gene-taxa correlations were visualized using corrplots [38] in R, where the strength of the correlation is indicated by the color and size of the visualization element (square) and the significance of the correlation is indicated via asterisk. We also computed the Sparse Correlation for Compositional Data (SparCC) [39] for the taxa found significantly correlated (q-value < 0.1) with the CRC genes. Pseudo p-values were computed using 100 randomized sets. Significant gene-microbe correlations (q-value < 0.1) and significant microbe-microbe correlations (SparCC |R| >=0.1 and p-value <0.05) were visualized as a network using Cytoscape v3.5.1 [40].

## Results

### Host RNA-Seq sample preprocessing and quality assessment

We first examined gene expression in colonic biopsies from 18 CF and 15 healthy individuals. Overall, CF and healthy samples had comparable number of reads (28,250,473 and 30,041,827 reads on average, respectively) with the average quality greater than 30 phred score across all samples (Fig S2). The sequences were annotated to generate estimated read counts and transcripts per kilobase million (TPM) using kallisto, [25] resulting in 173,259 total transcripts, of which 56,283 passed the filter of mean TPM greater than 1 (TPM>1). While the Principal component analysis (PCA) plots showed an overlap between the expression profile of most samples from CF and healthy individuals, it identified two possible outliers (samples 1096 and 1117) (Fig S3). In addition, the top five transcripts driving the PC were of mitochondrial origin (Fig S4). Hence, to reduce any bias in identifying differentially expressed genes, we filtered out all the mitochondrial transcripts from the data. We further investigated the outliers using the remaining transcripts by calculating cook’s distance between the samples, and found that the two samples (1096 and 1117) were still outliers (Fig S5). This was further evident by the heatmap of the top 20 most highly expressed genes (Fig S6), where we found an alternate expression pattern for the two samples, compared to the rest. Therefore, the two outlier CF samples (1096 and 1117) were eliminated from further analysis.

### Differentially expressed host genes between CF and healthy mucosal samples

To examine gene expression differences we used read counts from the remaining 16 CF and 15 healthy samples. Using DESeq2 we identified 1543 differentially expressed genes at q-value < 0.05 (Benjamini-Hochberg correction; see Fig S8 for a volcano plot). Of the 1543 differentially expressed genes, 919 (59%) were up-regulated and 624 (41%) were down-regulated in CF patients. Including sex as a covariate in the model did not susbstantially alter the results (only 43 additional differentially expressed genes were identified); therefore, we did not include sex in downstrem analyses. The full list of differentially expressed genes significant at q-value < 0.05 is available in Additional File 2.

We visualized the expression pattern of six representative genes, selected from genes included in the colorectal cancer disease pathway (Figure 1A). Consistent with the expectation of changes in mucosal immunity that could compensate for a diminished protective mucus function, we noted *LCN2* to be one of the top differentially expressed genes (q-value = 2.54E-08, Wald test). *LCN2* encodes for lipocalin 2, which limits bacterial growth by sequestering iron-laden bacterial siderophore [41]. However, a number of other top genes are involved in major cellular biology processes, and were previously related to cancer pathogenesis and colon cancer. Examples include *RRS1* (q-value = 6.16E-09), which encodes for the ribosomal biogenesis protein homolog that promotes angiogenesis and cellular proliferation, but suppresses apoptosis [42]; *KRTAP5-5* (q-value = 4.89E-08), which encodes for keratin-associated protein 5-5, a protein that plays important roles in cytoskeletal function and facilitates various malignant behaviors that include cellular motility and vascular invasion [43]; *ALDOB* (q-value = 2.64E-07), which encodes for aldolase B, an enzyme that promotes metastatic cancer-associated metabolic reprogramming [44]. Additional examples of differentially expressed genes (Log-fold change > 0.5 and q-value < 0.05), such as *CDH3, TP53INP2, E2F1, CCND2, and SERPINE1,* were also previously shown to have direct roles in colorectal and digestive cancers [45–47]. While, some of these genes participate in basic cancer-related cellular functions such as proliferation and invasion [45–52], others, e.g. *BEST2,* play important roles in gut barrier function and anion transport [53,54]. In addition to the genes visualized in Figure 1A, additional randomly selected differentially expressed genes are visualized in Additional File 1 (Fig S10), showing expression pattern difference between the CF and healthy samples.

**Figure 1.**
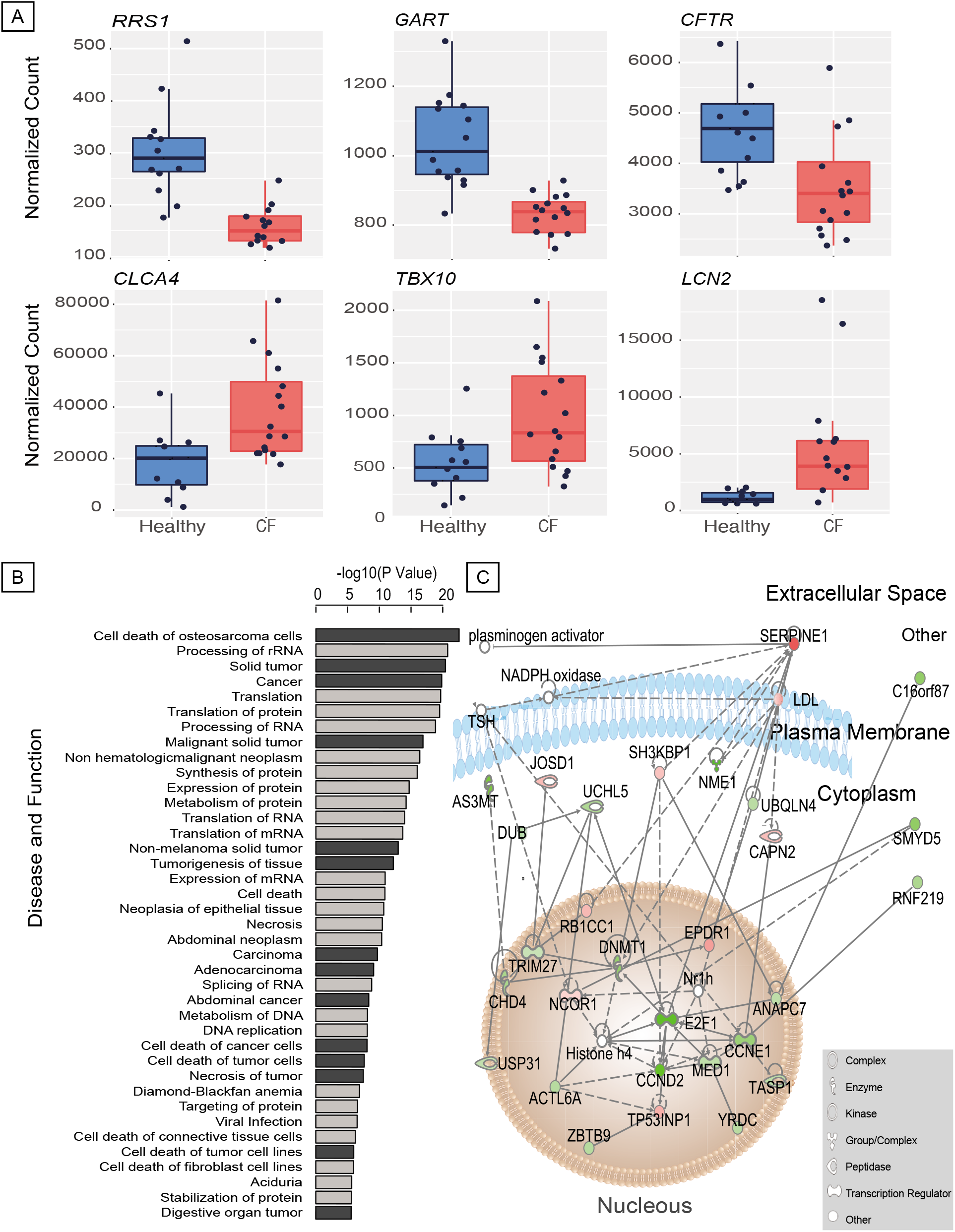
Differentially expressed (DE) host genes between CF patients and healthy individuals. (A) Box plots showing the expression level of six significantly DE genes that are a part of the gastrointestinal cancer pathway (B) Functional and disease categories that are enriched among DE genes sorted by the p-value (y-axis)), with darker bars indicating cancer-related categories. (C) Gene-gene interaction network showing genes in the gastrointestinal cancer pathway with green representing up-regulated genes in CF and red representing down-regulated genes in CF. The intensity of the color is indicative of more (brighter) extreme or less (duller) log-fold change measurement in the dataset. The shapes represent the protein function, illustrated in the legend at the bottom right. The figure is structured to show the cellular location in which each gene is active.

We next performed an enrichment analysis to categorize functional and disease pathways among differentially expressed genes (q-value <0.05) in IPA. The top canonical pathways (Fig S11) are mostly responsible for signaling and regulatory functions, such as *EIF2* signaling (p-value = 3.32E-35), mTOR signaling (p-value = 3.83E-08) and regulation of chromosomal replication (p-value = 1.60E-06). Of the 39 significantly enriched disease and functional pathways (p-value < 1.00E-05; Figure 1B), 14 are related to cancer, including gastrointestinal cancer (p-value = 2.61E-06), abdominal cancer (p-value = 9.23E-03), large intestine cancer (p-value = 7.00E-05), and colorectal cancer (p-value = 8.63E-03). In addition, using the list of differentially expressed genes we found that the promoter sequences are enriched with binding sites of 96 potential transcription regulators (p-value < 0.01; see Methods). Among these transcription factors, many have been previously shown to control cancer related pathways. For example, *MYCN* and *KRAS* are prominently involved in neuroblastoma and colorectal cancer, respectively [55,56]. *NHF4A* is involved in transcriptional regulation of many aspects of epithelial cell morphogenesis and function, which has been linked to colorectal cancer [57]. *CST5,* which encodes cytostatin D, is a direct target of p53 and vitamin D receptor, promotes mesenchymal-epithelial transition to suppress tumor progression and metastasis [58]. *E2F3* is a potent regulator of the cell cycle and apoptosis that is commonly deregulated in oncogenesis [59,60].

A metabolic network for the gastrointestinal (GI) cancer-related differentially expressed genes is shown in **Figure 1C**, illustrating the interactions between genes that are up-regulated in CF (Eg. *TP53INP1, SERPINE1, NCOR1* and *CAPN2)* and down-regulated in CF *(E2F1, MED1, ECND2 and AS3MT),* highlighting the cellular location of these genes’ product. Additional gene network for colorectal cancer can be found in Additional file 1 (Fig S12), where the genes are also positioned in the region of the cell where they are most active. We found that genes such as *BEST2* (involved in ion transport) and *RUVBL1* (involved in cell cycle, cell division, and cell damage) are down-regulated, while genes such as *TP53INP2* (involved in transcription regulation) and *CDH3* (involved in sensory transduction) are up-regulated. Given the predicted role of gene regulation in colorectal cancer and the dysregulation of CRC-related pathways, these results may help understand mechanisms controlling early onset of colon cancer in cystic fibrosis.

### Difference in microbiome composition between CF and healthy gut mucosa

To further understand the potential of altered microbiota-host interaction in the CF colon, we next investigated differences in the composition of the mucosal microbiome between CF and healthy individuals. We found a significant difference between beta-diversity of gut mucosal microbiome in CF patients compared to healthy individuals with respect to unweighted UniFrac and non-phylogenetic Bray-Curtis metrics (Adonis p-value = 0.001). As observed in the PCoA plot (Figure 2A) the samples were clustered based on their disease condition (CF or healthy). The overall biodiversity of mucosal microbiome was depleted in CF compared to healthy samples, which is depicted by significant decrease in alpha diversity measured by Chao1 (p-value = 0.015, Wilcoxon rank sum test, Figure 2A) and Observed OTUs (p-value = 0.024, Wilcoxon rank sum test, in Additional File 1 (Fig S13)) metrics in CF relative to healthy controls.

**Figure 2.**
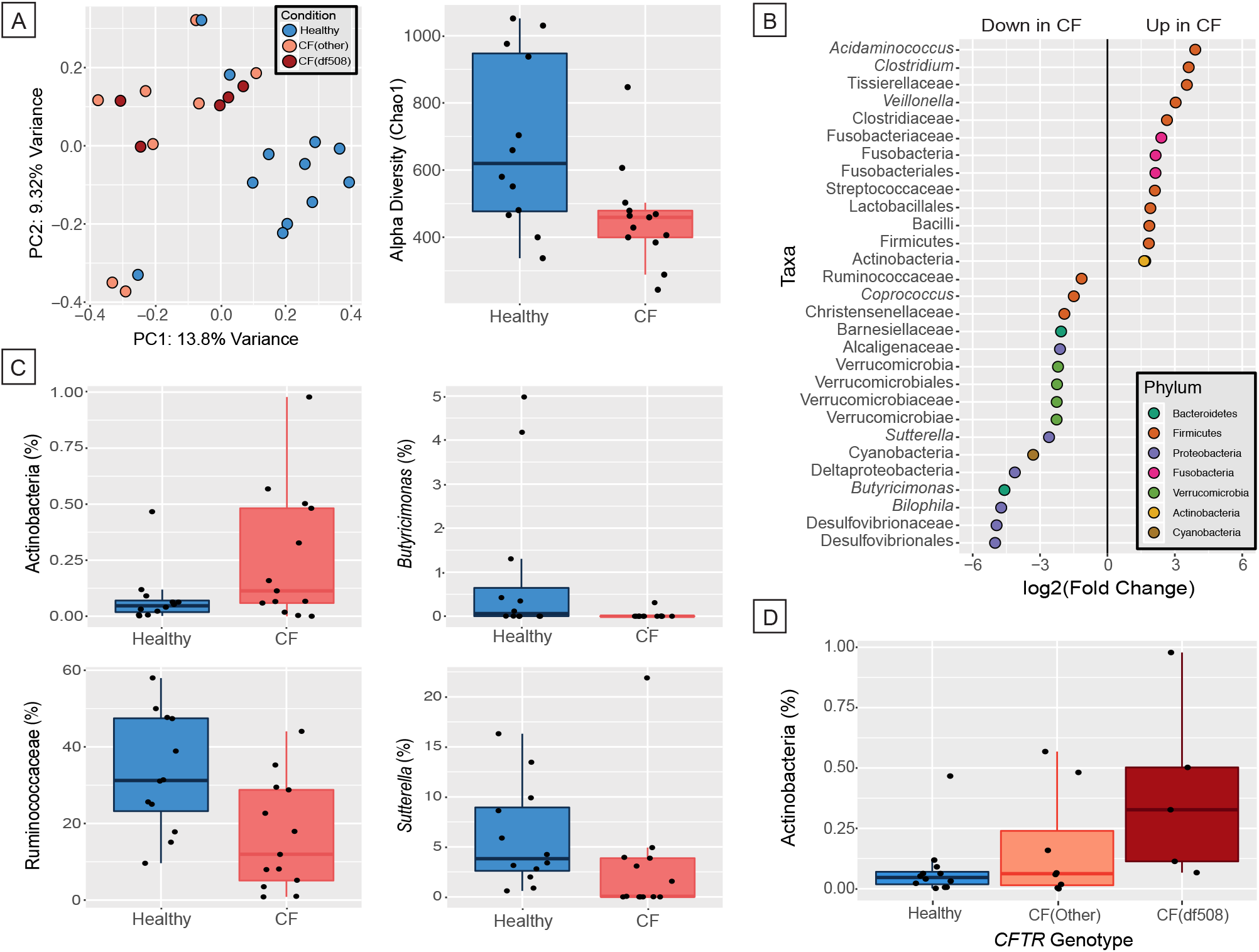
Differences between CF and healthy gut mucosal microbiota. (A) (left) Principal coordinate analysis plot based on Bray-Curtis distance indicating difference in beta-diversity between CF and healthy gut mucosal microbiome. The axes represent the percentage variance along the first two principal components, and the color of samples indicates their mutation status, i.e. healthy (i.e. no known mutation in CFTR), CF(df508) (homozygous for the DF508 mutation), and CF(other) (either one or zero alleles of the DF508 mutation); (right) Boxplot depicting difference in alpha-diversity for Chao1 metric between CF and healthy gut microbiome. (B) Dotplot showing significantly differentially abundant taxa (q-value < 0.1) between CF and healthy samples. The taxa are listed along the y-axis and are colored by their phylum, and the x-axis indicates the log2 fold-change in CF compared to healthy as baseline. (C) Boxplots indicating the percentage relative abundance of taxa showing differential abundance between CF and healthy gut microbiome (q-value < 0.1). (D) Boxplot depicting the abundance of Actinobacteria for three mutation levels – Healthy, CF(other) and CF(df508).

We assessed the changes in abundance of microbes at various taxonomic levels between CF and healthy gut mucosal microbiome using phyloseq. We found 51 OTUs that were significantly differentially abundant between CF and healthy individuals (q-value < 0.1, Additional file 3). At different taxonomic ranks, we found 7 genera, 10 families, 4 orders, 4 classes, and 5 phyla differentially abundant between CF and healthy samples (q-value < 0.1 by Wald’s test; Additional file 3). Overall, an increased abundance in taxa, predominantly belonging to Firmicutes and Fusobacteria, was observed in CF individuals compared to healthy controls, while taxa belonging to Bacteroidetes, Verrucomicrobia, and Proteobacteria phyla showed a marked decrease in patients with CF relative to healthy controls (Figure 2B). In particular, there was an increase in abundance of class Actinobacteria in individuals with CF compared to healthy controls (q-value = 0.079), while *Butyricimonas* (q-value = 0.009), Ruminococcaceae (q-value = 0.081), *Sutterella* (q-value=0.040) were found depleted in CF samples (Figure 2C). Additional examples of differentially abundant taxa between CF and healthy samples can be found in the Additional file 1 (Fig S14).

Next, we tested whether *CFTR* genotype, which affects disease severity, is associated with variation in the microbiome. Specifically, we hypothesized that variation in the microbiome is correlated with the number of alleles of the DF508 mutation, a deletion of an entire codon within *CFTR* that is the most common cause for CF. To test this, we performed likelihood ratio test to identify differentially abundant taxa between three genotype classes: CF-DF508 (homozygous for the DF508 mutation), CF-other (either one or zero copies of the DF508 mutation), and healthy (no known mutations in *CFTR*). We found a gradient-like trend in abundance for Actinobacteria (q-value = 0.081), showing increase in abundance with increasing severity of mutation status (Figure 2D).

To assess the potential functional changes in the microbiome, we predicted abundance of metabolic pathways and enzymes using the PICRUSt pipeline [36] and KEGG database, and compared them for differences between CF and healthy individuals. Seven predicted pathways (as defined by KEGG level 3) were found to be differentially abundant between CF and healthy: bacterial toxins were enriched in CF compared to healthy, while propanoate metabolism, restriction enzyme, pantothenate and CoA biosynthesis, thiamine metabolism, amino acid related enzymes, and aminoacyl-tRNA biosynthesis were depleted in CF compared to healthy (q-value < 0.2 using Wilcoxon rank sum test; in Additional File 1 (Fig S15)).

### Interactions between gastrointestinal cancer-related host genes and gut microbes

In order to investigate the relationship between host genes and microbes in the colonic mucosa and their potential role in the pathogenesis of gastrointestinal cancers in CF patients, we considered correlations between 250 differentially expressed genes enriched for GI cancers and 35 microbial taxa (collapsed at genus or last characterized level, and filtered at 0.1% relative abundance, see Methods). Using Spearman correlations, we found 50 significant unique gene-microbe correlations in the gut (q-value < 0.1), where the magnitude of correlation (Spearman rho) ranged between (−0.77, 0.79) (Additional file 4). Interestingly, most of the taxa that significantly correlated with the genes also differed significantly in abundance between CF and healthy individuals. We visualized all the correlations between taxa abundance and host gene expression in **Figure 3A**. In particular, we found some significant positive gene-taxa correlations (q-value < 0.05), between *Butyricimonas* and *ZNHIT6* (Spearman rho = 0.76), Christensenellaceae and *MDN1* (Spearman rho = 0.78), and *Oscillospira* and *NUDT14* (Spearman rho = 0.79). A few significant negative correlations (q-value < 0.05), such as between Christensenellaceae and *TBX10* (Spearman rho = – 0.78), and Ruminococcaceae and *LCN2* (Spearman rho = −0.77) were also found.

**Figure 3.**
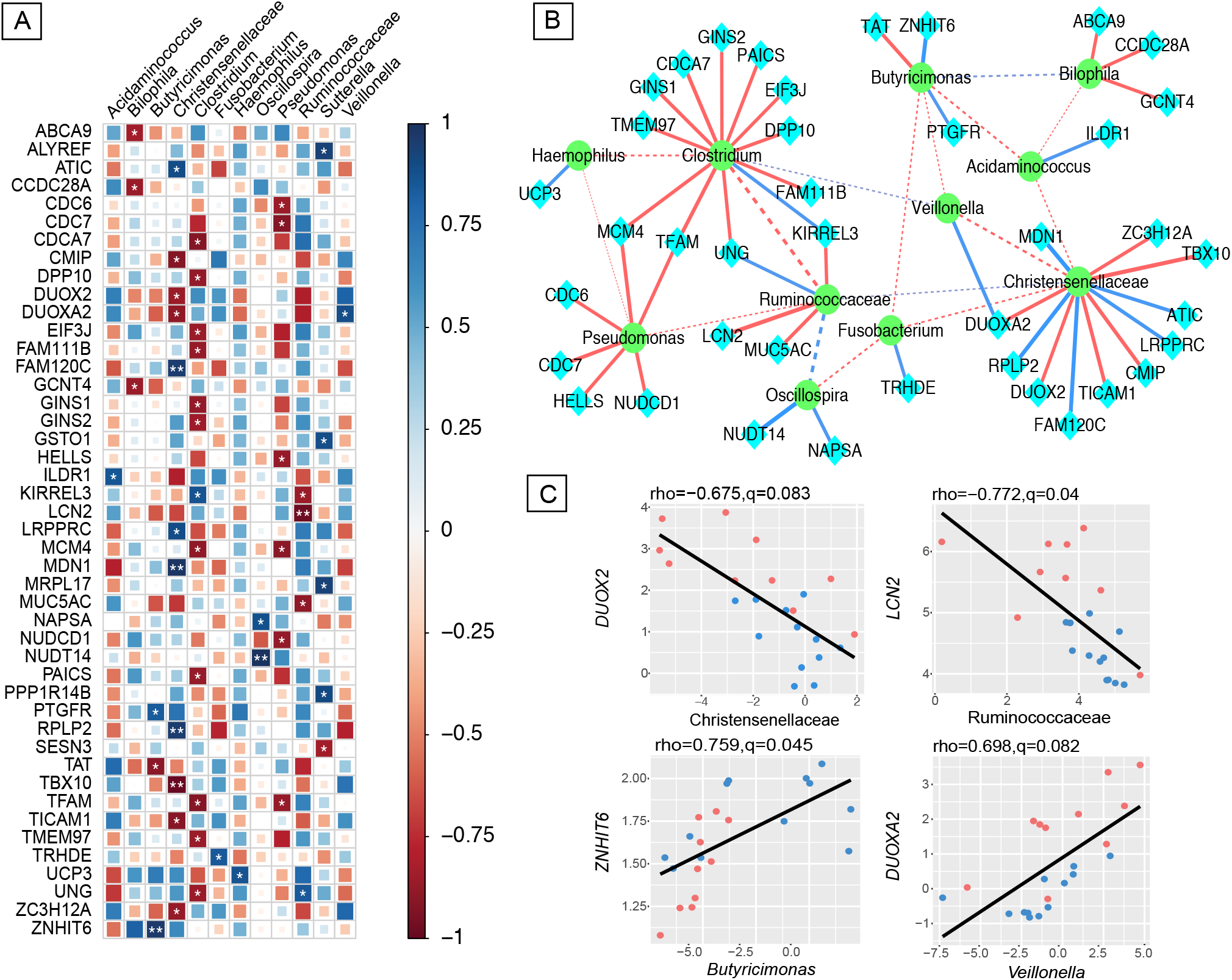
Interactions between gastrointestinal cancer-related host genes and gut mucosal microbes. (A) Correlation plot depicting gene-microbe correlations. Color and size of the squares indicate the magnitude of the correlation, asterisks indicate significance of correlation (** indicates q-value < 0.05 and * indicates q-value < 0.1). (B) Network visualizing the significant gene-microbe correlations (solid edges, q-value < 0.1) and significant microbe-microbe correlations (dashed edges, SparCC |R| >=0.1 and p-value < 0.05). Blue edges indicate positive correlation and red edges indicate negative correlation. Edge thickness represents the strength of the correlation. (C) Scatterplots depicting pattern of grouping by CF (red) and healthy (blue) samples in a few representative gene-microbe correlations, where the strength of correlation (Spearman rho) and significance (q) are indicated on the top.

To characterize potential microbe-microbe interactions in our dataset, we computed correlations between the microbes significantly correlated (q-value <0.1) with the genes using sparCC (see Methods and Additional file 4) [39]. The notable aspects of the significant gene-microbe correlations (q-value < 0.1) and significant microbe-microbe correlations (SparCC |R| >=0.1 and pseudo-P value <0.05) are graphically represented in **Figure 3B**, where solid edges denote gene-microbe correlations and dashed edges represent microbe-microbe correlations. This subnetwork of microbe-microbe correlations depicts correlated abundance changes in the microbiome as a function of their presence (**Figure 3B**, dashed edges). For instance, *Bilophila* and *Butyricimonas* are both depleted in CF (q-value < 0.05), and the abundance of the two genera is also correlated across individuals (SparCC R = 0.5, pseudo-P value = 0.04). On the other hand, Ruminococcaceae was found depleted in CF (q-value = 0.081), while *Clostridium* was enriched in CF (q-value = 0.0004), and this inverse co-occurrence pattern leads to a negative correlation between the two taxa across study participants (SparCC R = – 0.66, pseudo-P value = 0). Furthermore, in the gene-microbe subnetwork (**Figure 3B**, solid edges), microbial nodes have more edges on average compared to genes, where Christensenellaceae and *Clostridium* formed distinct hubs in the network. This potentially implies that these microbes and their pathways are shared across multiple GI cancer-associated genes. Of note, *Bilophila, Clostridium,* and *Pseudomonas* are mostly negatively correlated with GI cancer genes, while *Haemophilus, Oscillospira, Veillonella, Fusobacterium and Acidaminococcus* are only positively correlated with GI cancer genes (q-value < 0.1).

In addition to the overall network, **Figure 3C** depicts pairwise correlations between host gene expression and microbial taxa where both have been previously linked to CRC, and thus may be of interest. For example, *LCN2*, known to be overexpressed in human CRC and other cancers (Maier et al. 2014), is negatively correlated with Ruminococcaceae (Spearman rho = −0.77, q-value = 0.040), which is found depleted in CRC [61,62]. Both *DUOX2* and *DUOXA2* are found to be negatively correlated with Christensenellaceae (Spearman rho < −0.65, q-value < 0.1), while *DUOXA2* is positively correlated with *Veillonella* (Spearman rho = 0.70, q-value = 0.082). *DUOX2* and its maturation factor *DUOXA2* are responsible for H2O2 production in human colon and are known to be up-regulated in gastrointestinal inflammation [63,64]. Christensenellaceae, a heritable taxon [65], has been shown to decrease in abundance in conventional adenoma [62], a precursor of CRC, whereas *Veillonella,* which is known to be pro-inflammatory, is found to be represented in human CRC [66]. Thus, the pattern of grouping by CF and healthy samples in these representative correlations are found to be similar to known associations in CRC and other gastrointestinal malignancies.

## Discussion

Recent advances in the treatment have significantly prolonged the lives of CF patients [67,68]. However, this led to new challenges, such as an elevated risk for gastrointestinal cancer [2,69]. Thus, CF patients show 5-10-fold increased risk of CRC compared to healthy individuals, and that increases even further with immunosuppressive drugs [3,6]. Understanding the molecular mechanisms that control the increased risk is key for early detection and the development of tailored treatments[6]. The importance of interactions between host and microbiome in the pathogenesis of colorectal cancer has become increasingly clear [61,70,71]. To understand the role of these interactions in CF, we jointly profiled host colon gene expression and mucosal microbiome composition data in CF patients and healthy controls. We observed an enrichment of cancer-associated dysregulated genes — specifically colon cancer — in CF patients compared to healthy controls. We also observed a shift in the microbiome and identified strains previously linked to colon cancer that varied in their abundance between CF and healthy individuals. We further found relevant correlations between these cancer enriched genes and microbes that may illuminate the mechanisms of CRC development in CF patients.

Several previous studies have studied the role of host gene regulation in CF patients [5,72,73]. While results from previous studies are based on either phenotypic observations, examining candidate genes such as *CFTR,* or an exploration of gene expression data from respiratory or blood samples [5,72,73], our work is the first, as far as we know, that focused on a comprehensive transcriptomic analysis of colon biopsies. This allowed us to characterize patterns of host gene regulation specific to the CF colon epithelium. In addition to an enrichment of cancer-related pathways among genes that are differentially expressed in CF, we also observed an enrichment for immune response pathways, including signal transduction, cell adhesion, and viral infection. Interestingly, one of the most significant pathways enriched in our current data, the eIF2 signaling pathway, has been previously shown to play an important role in immune response, and cells with defective eIF2 signaling pathway were more susceptible to bacterial infections [74]. In addition, our analysis revealed that tumor suppressor genes are differentially regulated in the colon of CF patients. In addition to *CFTR,* we found other tumor suppressor genes, such as *HPGD,* to be down-regulated in CF patients colon. *HPGD* was previously shown to be down-regulated in lungs of CF patients [5,75]. Down-regulation of these tumor suppressor genes can lead to predisposition of colon cancer [42,76,77], suggesting a potential mechanism underlying the reported increased risk and early development of colon cancer in CF patients [5,69].

In addition to host gene regulation, the microbiome has also been implicated in the development of many diseases, including CRC [61,78]. In the context of CF, previous studies have focused on characterizing shifts in the fecal or airway microbiome [20,79]. Here, we profiled the colonic mucosal microbiome, with the goal of understanding its role in the development of CRC in CF patients. We found a clear distinction between microbiome populations from CF compared to healthy mucosa. Overall, similar to several other GI diseases, we also observed a reduced microbial biodiversity in the CF population [80]. We also found a depletion in butyrate producing bacteria, such as Ruminococcaceae and *Butyricimonas,* similar to previously reported depletion in butyrate producing microbes by Manor et al [20] in their study comparing CF fecal samples from children on varying degree of fat intake. Butyrate helps promote growth and can also act as an anti-inflammatory agent, and is therefore an important compound for colon health [20]. Interestingly, mice with compromised GI defense system also had a reduced number of butyrate producing bacteria, similar to our observations in the CF patients, who generally consume a high fat diet [81]. In addition to decrease in Ruminococcaceae, we also observed a depletion in another butyrate producing bacteria *Sutterella;* loss of abundance in both of these strains have been previously observed in CRC [61,82]. We also found an increase in Actinobacteria, one of the most predominant genera found in the sputum of CF patients [79,83], but decreased in colon cancer gut microbiome [84]. Furthermore, our observation of significant decrease in the abundance of Verrucomicrobia, and increase in abundance of Firmicutes and Actinobacteria in CF patients, is consistent with findings from the fecal microbiome of CF patients [21]. Lastly, we found an increase in predicted bacterial toxins in the CF population, which might be explained by the increase in pathogenic bacteria such as *Pseudomonas* and *Veillonella.* This can potentially damage epithelial cells or induce mutations leading to unfavorable clinical outcome [85,86].

Integrating mucosal microbiome and host gene expression profiles, we observed several correlations between differentially expressed colon epithelial genes and gut mucosal bacteria in CF. Co-culture and obligate cross-feeding studies have shown an increased virulence of a pathogen in the presence of other bacteria, thus triggering an immune response that can determine the clinical outcome [87,88]. One such example is the increased virulence of *Pseudomonas* in presence of *Veillonella* as seen in mice tumor model resulting in host clinical deterioration [88]. Interestingly, we found both of these microbes *(Veillonella* and *Pseudomonas)* in higher abundance in CF patients. Furthermore, we also found a strong correlation between *Veillonella* and *DUOXA2,* a highly expressed gene causing inflammation in ulcerative colitis [89]. Another such correlation that we observed was between highly expressed *LNC2* gene, which plays a role in innate immunity and has been previously found to be up-regulated in human colon cancers [90], and depletion of Ruminococcaceae, a butyrate producing bacteria that helps maintain colon health [20].

Our study has several limitations. First, CF patients have a high burden of antibiotic exposure. Since antibiotics affects the gut microbiome [91–93], this may impact the differences we observe between CF and healthy mucosal microbiome. It is challenging to account for this potential bias; although matched healthy controls that are also on antibiotics can be used, the effects of long-term antibiotics usage may be impossible to match in a non-CF control population. In addition, although we report potential host gene-microbe and microbe-microbe interactions, our study focused on correlations, and causality is not inferred. Although studying causality in host-microbe is challenging in humans, future studies using animal or cell models can be useful to disentangle the direction of interaction [94].

## Conclusions

To summarize, we report an analysis of the mucosal microbiome and host gene expression in the gut of CF patients and healthy controls. We find down-regulation of tumor suppressor genes, as well as up-regulation of genes that play a role in immune response and cause inflammation. Furthermore, we observe a shift in microbiome with depletion in butyrate producing bacteria that may help maintain colon health and increase in pathogenic strains in individuals with CF. Lastly, our study provides a set of candidate interactions between gut microbes and host genes in the CF gut. Our work sheds light on the role of host-microbiome interactions and their relevance for the early development of CRC in CF patients. Our results can provide clinicians and researchers with biomarkers that may potentially serve as targets for stratifying risk of CRC in patients with CF.

## Abbreviations

CF: cystic fibrosis;
CRC: colorectal cancer;
GI: gastrointestinal;
FDR: false discovery rate;
OTU: operational taxonomic unit;
PICRUSt: Phylogenetic Investigation of Communities by Reconstruction of Unobserved States;
KEGG: Kyoto Encyclopedia of Genes and Genomes.

## Acknowledgements

We thank the members of the Blekhman Lab for helpful discussions. This work was carried out, in part, by resources provided by the Minnesota Supercomputing Institute. This work is supported by award MNP 17.26 from the Minnesota Partnership for Biotechnology and Medical Genomics (R.B.), a McKnight Land-Grant Professorship from the University of Minnesota (R.B.), the Chainbreaker Breakthrough Cancer Research Grant from the Masonic Cancer Center at the University of Minnesota (A.K. and R.B), With One Breath (A.K.), and the Cystic Fibrosis Foundation (D.E.N.). Research reported in this publication was supported by the National Center for Advancing Translational Sciences of the National Institutes of Health Award Number UL1-TR002494. The content is solely the responsibility of the authors and does not necessarily represent the official views of the National Institutes of Health.

