## Additional file 1 for "Interactions between the gut microbiome and host gene regulation in cystic fibrosis"

* these authors contributed equally

**Supplemental Figures:**

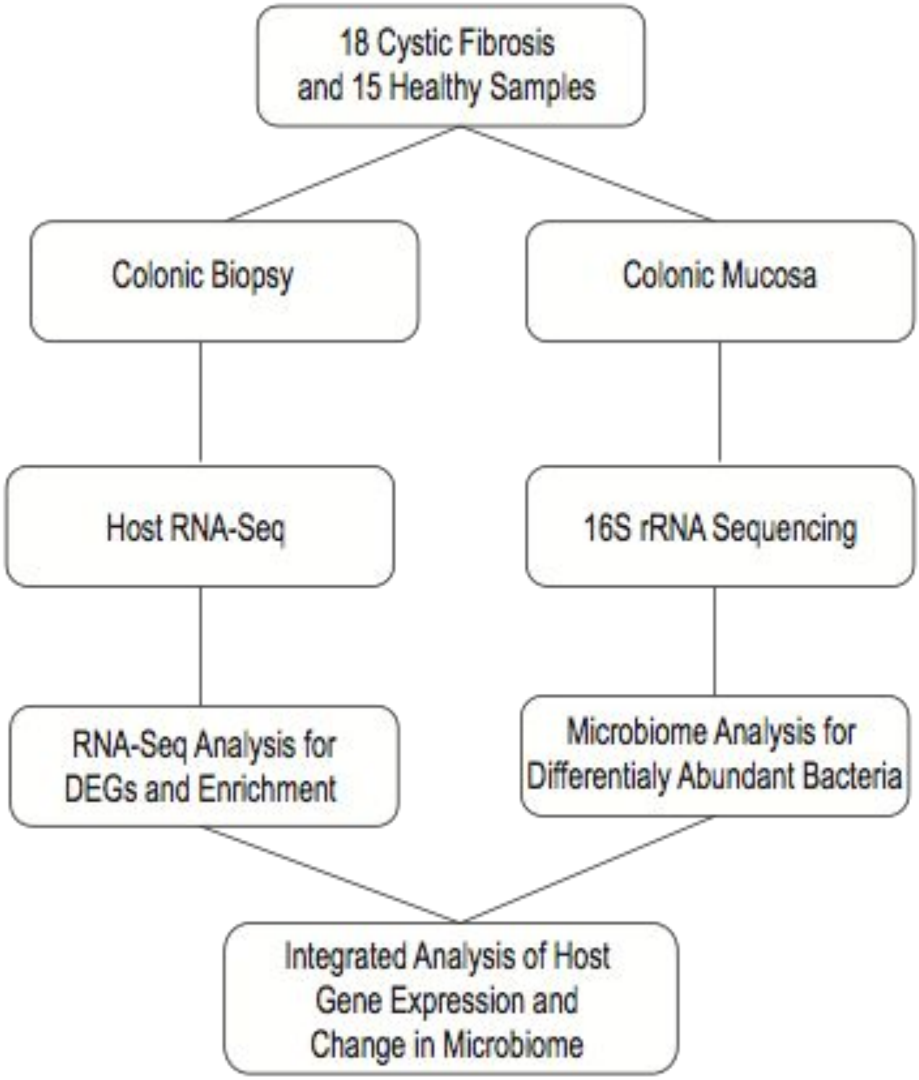


Fig S1. Experimental pipeline. Flowchart to illustrate major steps of the study, including sample information, host RNA-Seq analysis, microbiome analysis and integrated analysis.

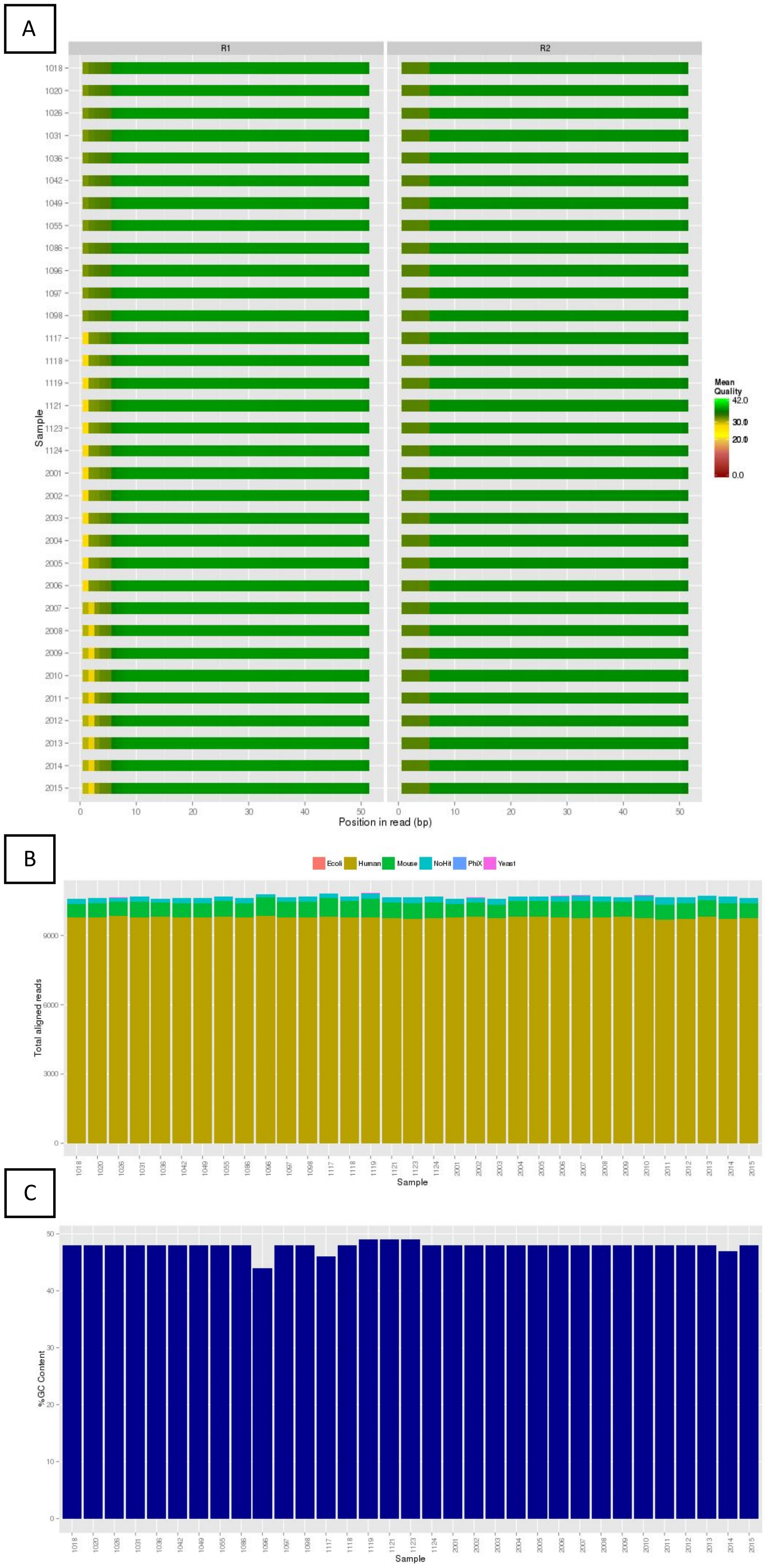


Fig S2. Quality control of RNA-seq data. (A) Average phred score of the forward (R1) and reverse (R2) reads, (B) Number of reads mapped to human and other species, (C) Total number of reads per sample

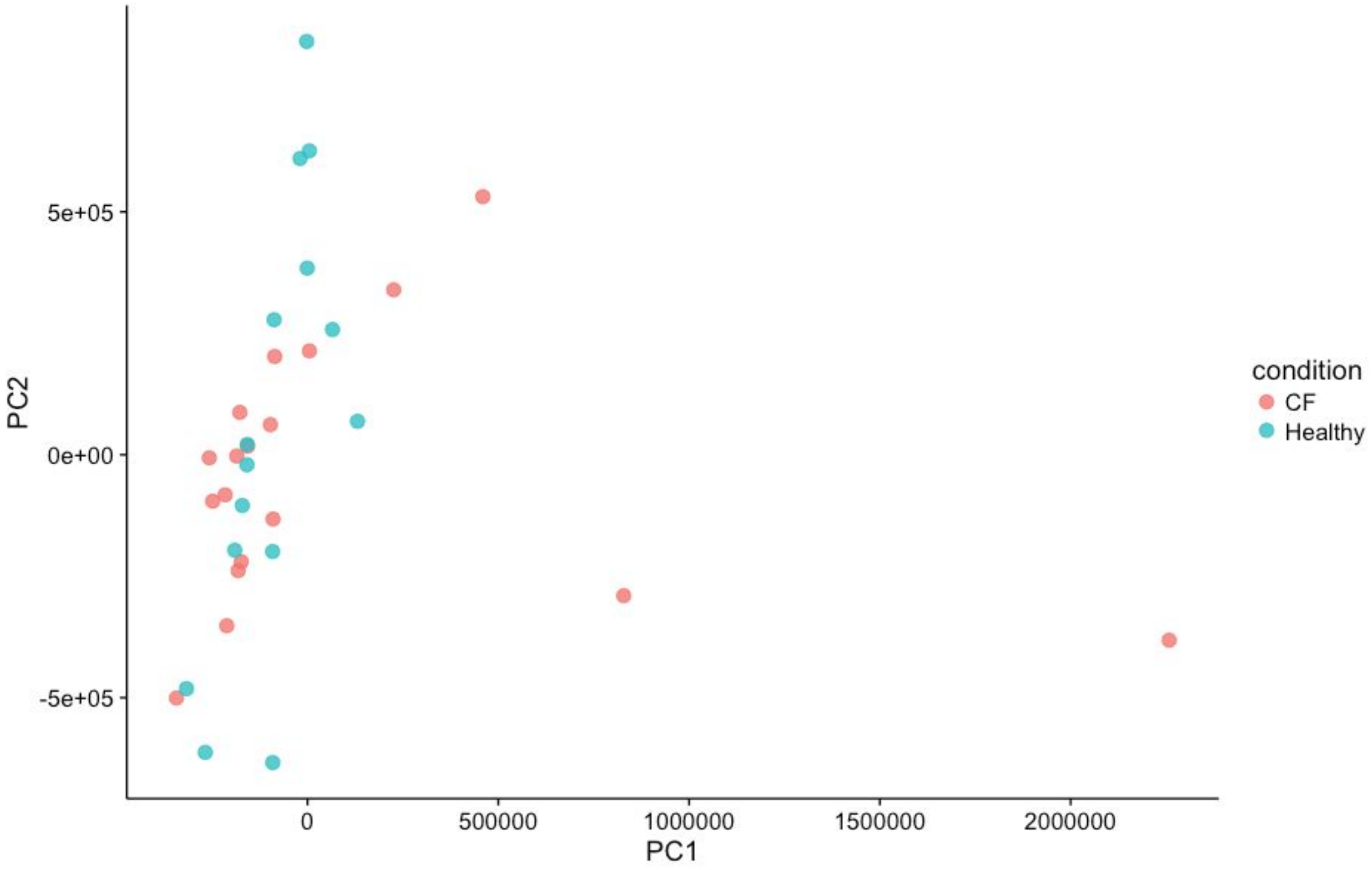


Fig S3. Sample clustering. PCA plot colored by condition, cystic fibrosis samples in red and healthy samples in green

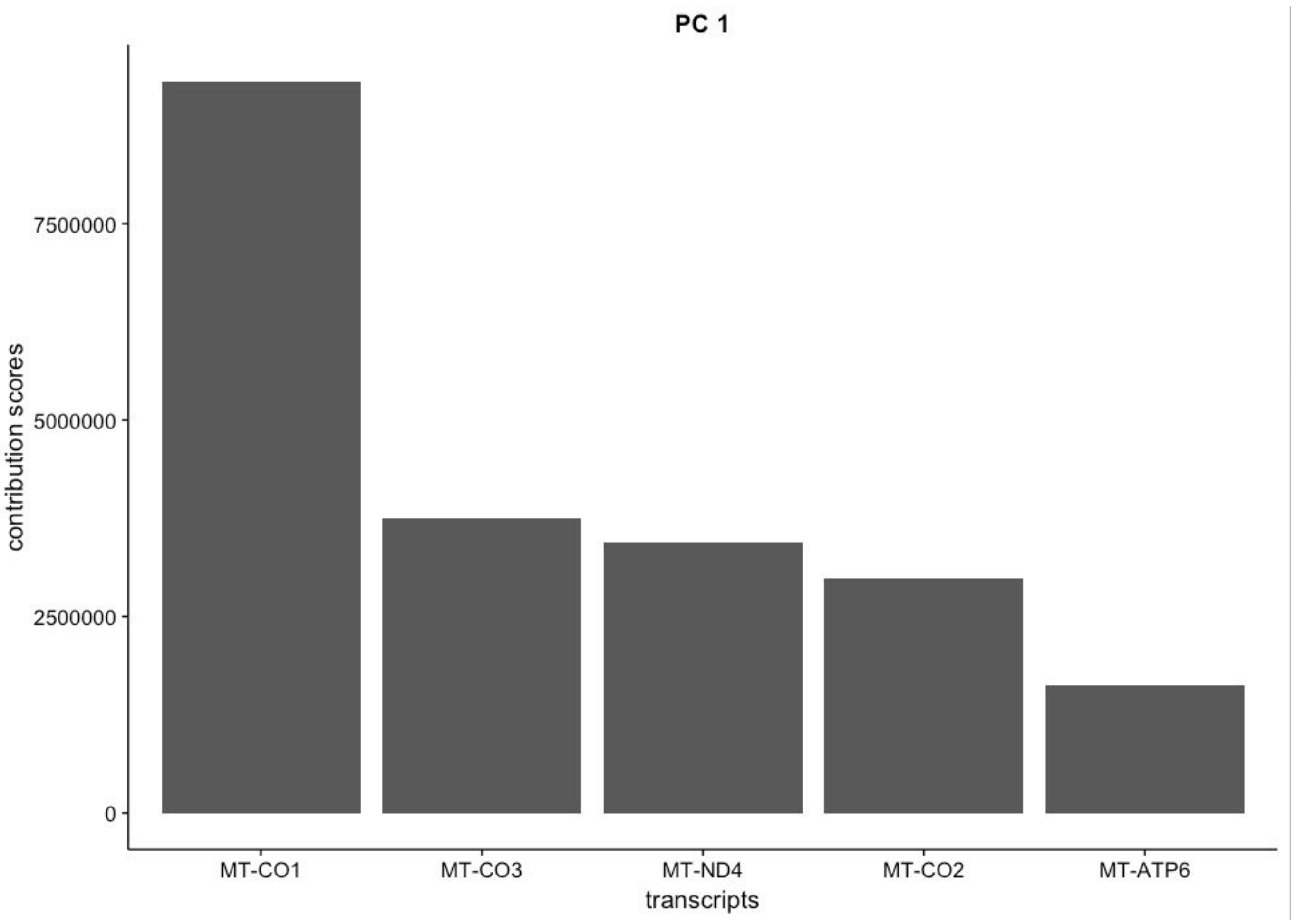


Fig S4. Primary genes defining the PC. Bar plots representing the top 5 genes driving PC1

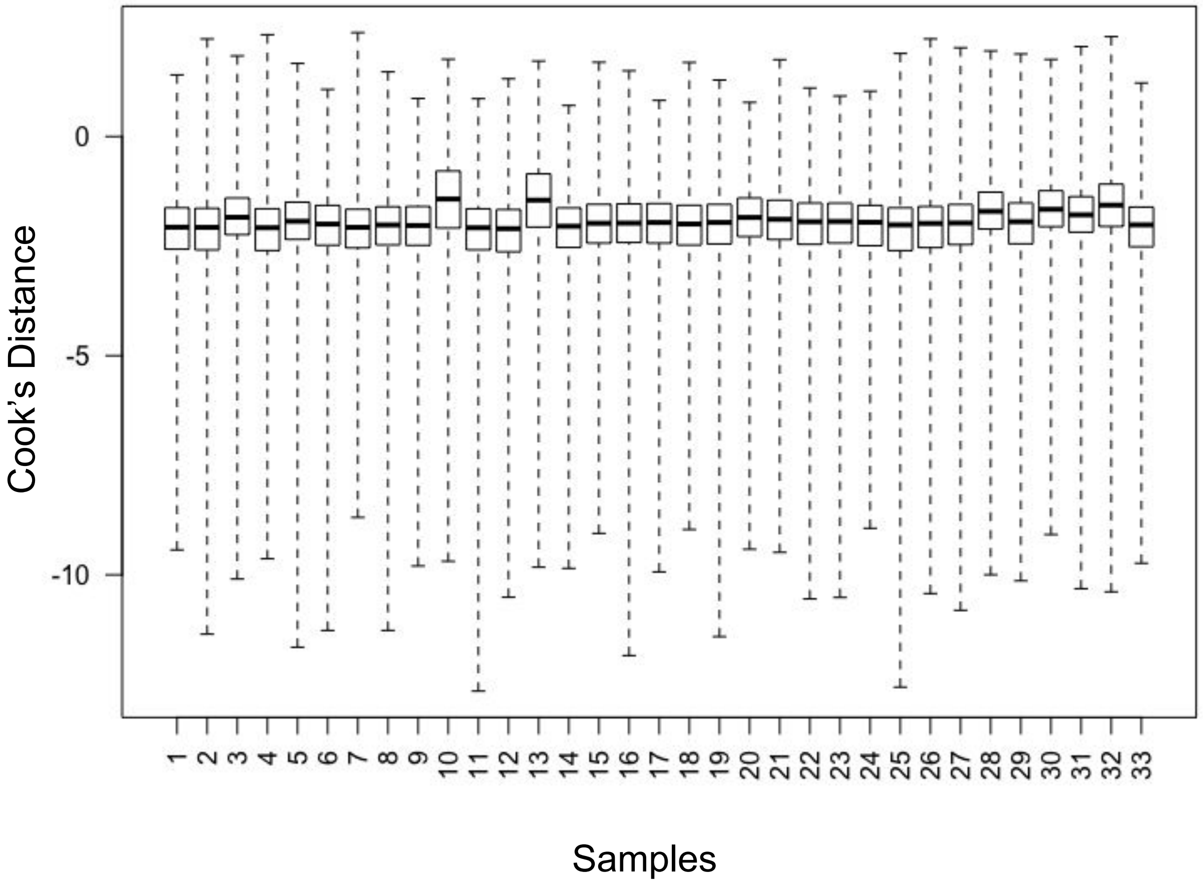


Fig S5. Quality control of samples. Box plots generated using cook's distance for determining any outlier samples

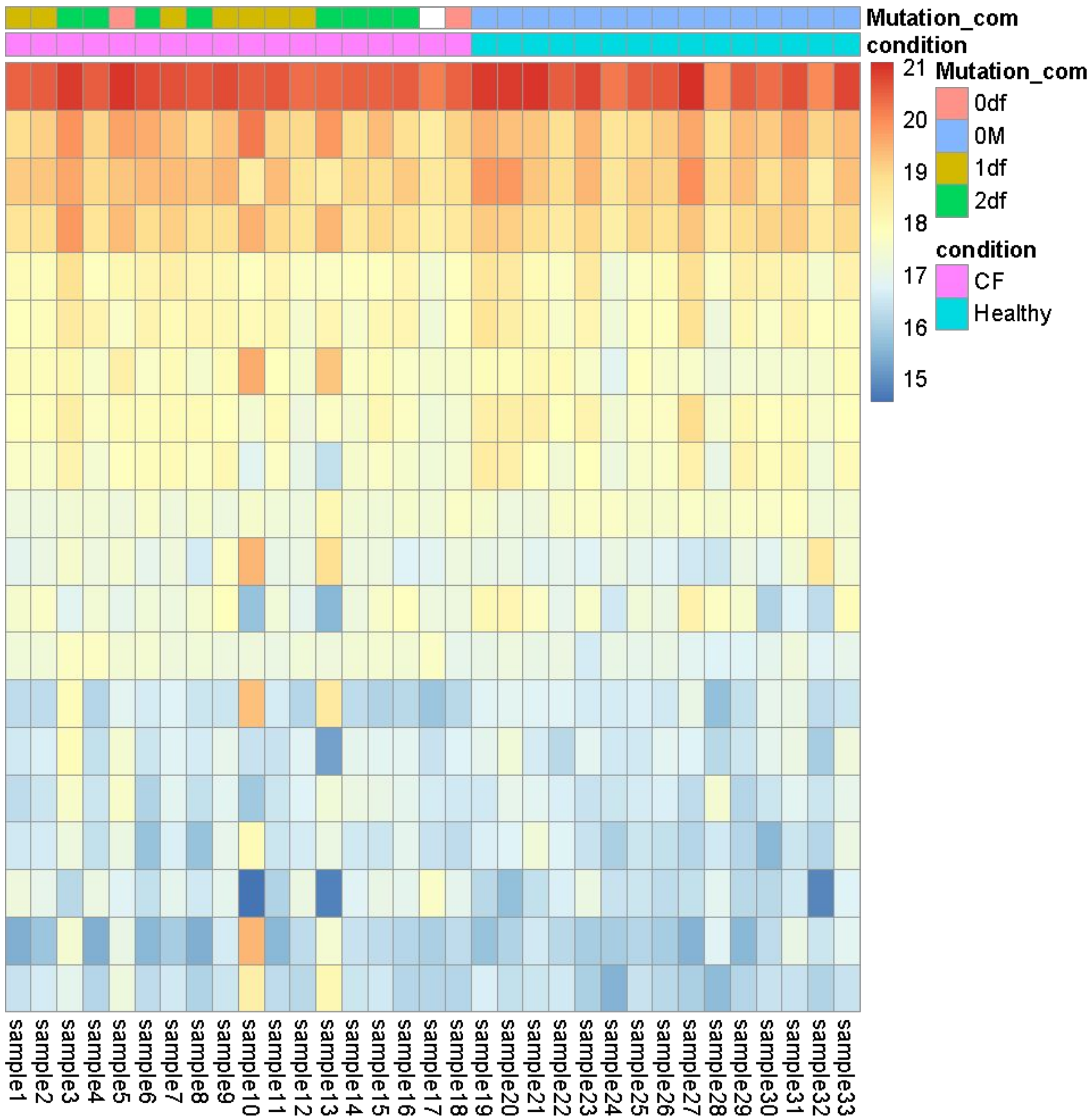


Fig S6. Transcript abundance. Heat map of top 20 genes, clustered by condition (cystic fibrosis represented by pink and healthy by cyan). Also includes the df508 mutation information on the CFTR gene (0M is for no mutation, 0df is when there is a mutation other than df508 deletion, 1df is for heterozygous df508 deletion and 2df is when both copies are deleted). Sample 17 did not have genotypic information.

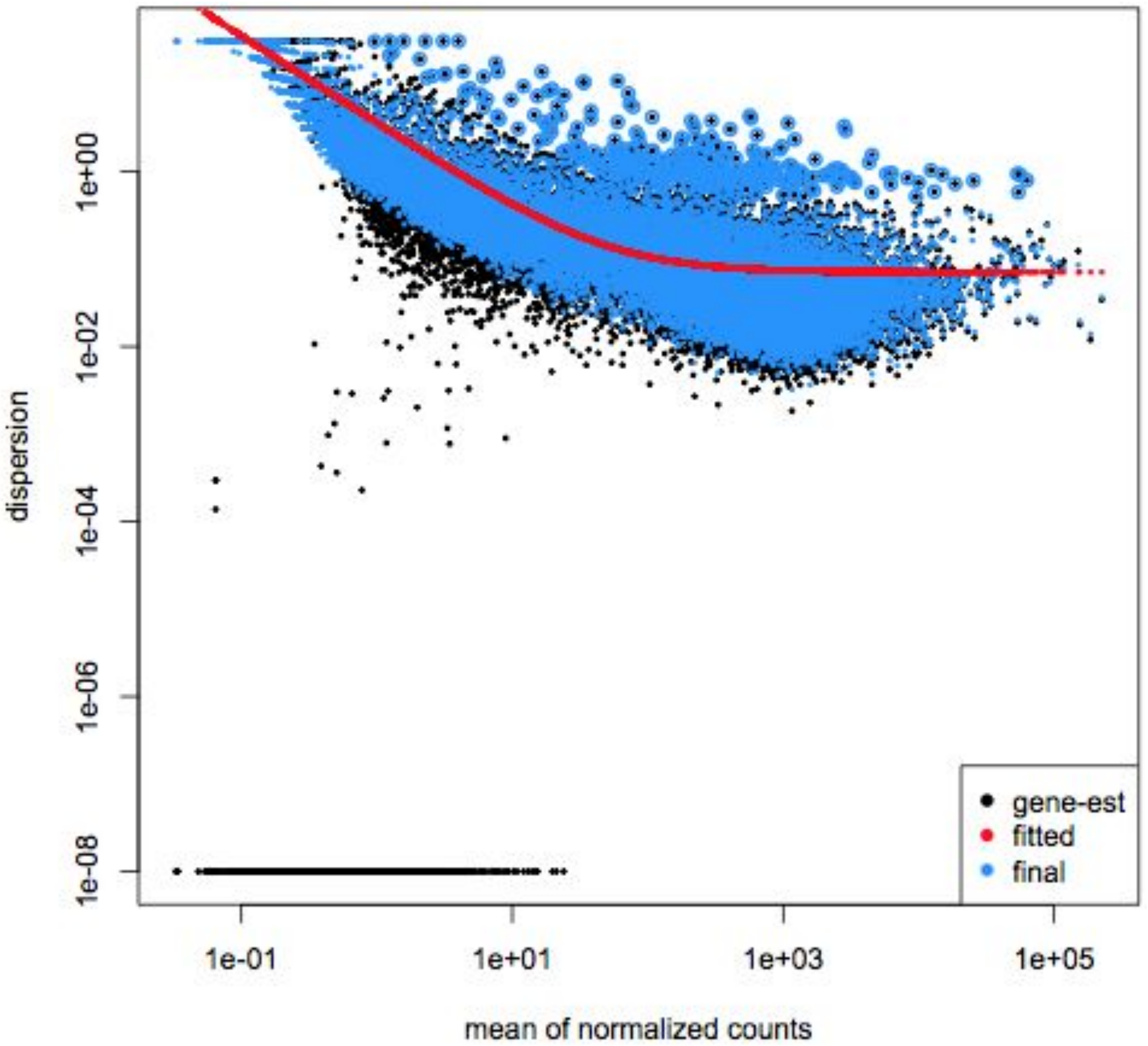


Fig S7. Data fitting. Dispersion plot for gene-wise estimates shrunken towards the fitted estimates (red line) and the outliers that were not shrunk are represented by the blue circles around the data points.

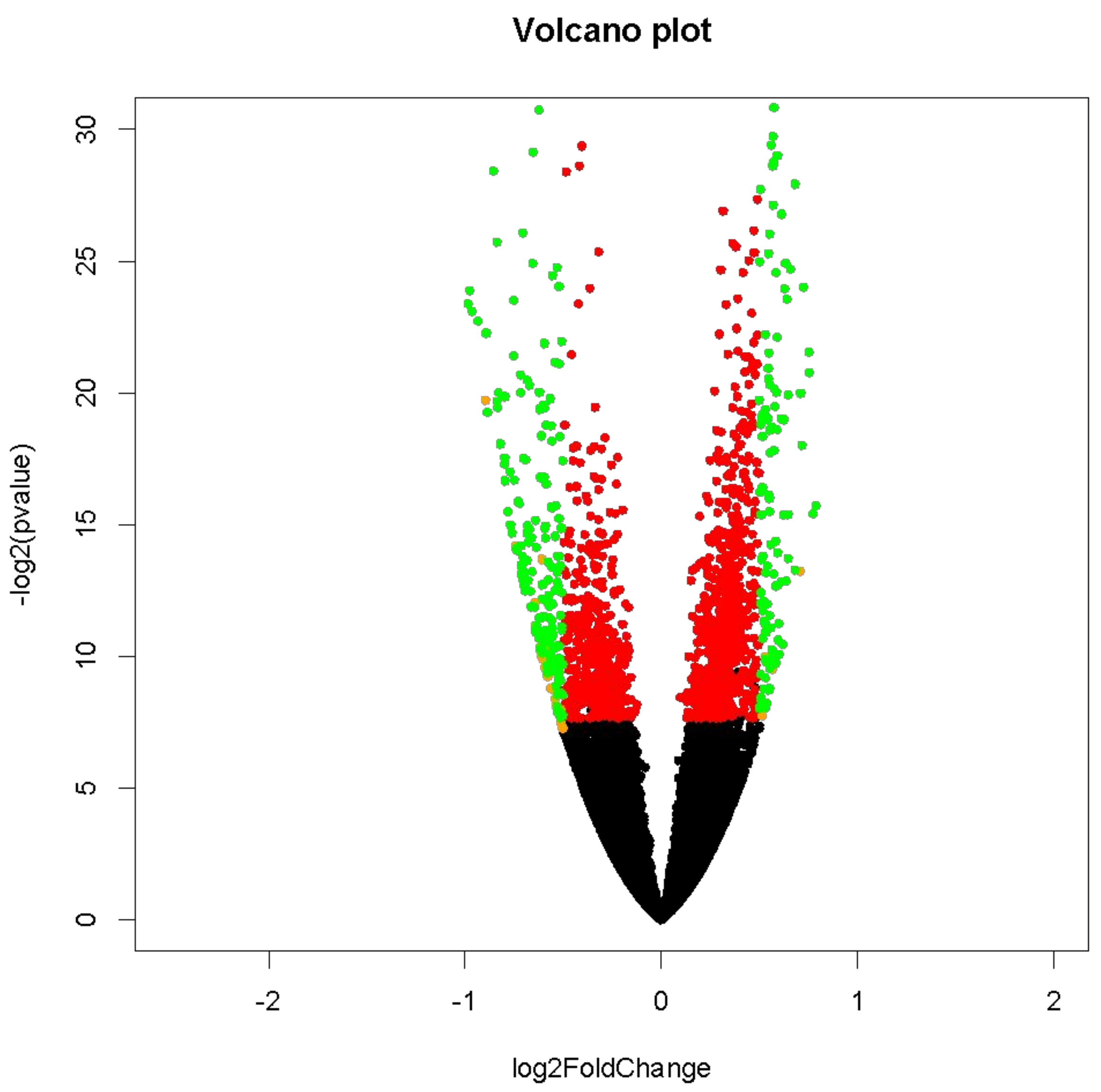


Fig S8. Differentially expressed genes (DEGs). Volcano plot showing all the DEGs, FDR < 0.05 represented by the red points and log fold change (log2|FC|) > 0.5 represented by green points.

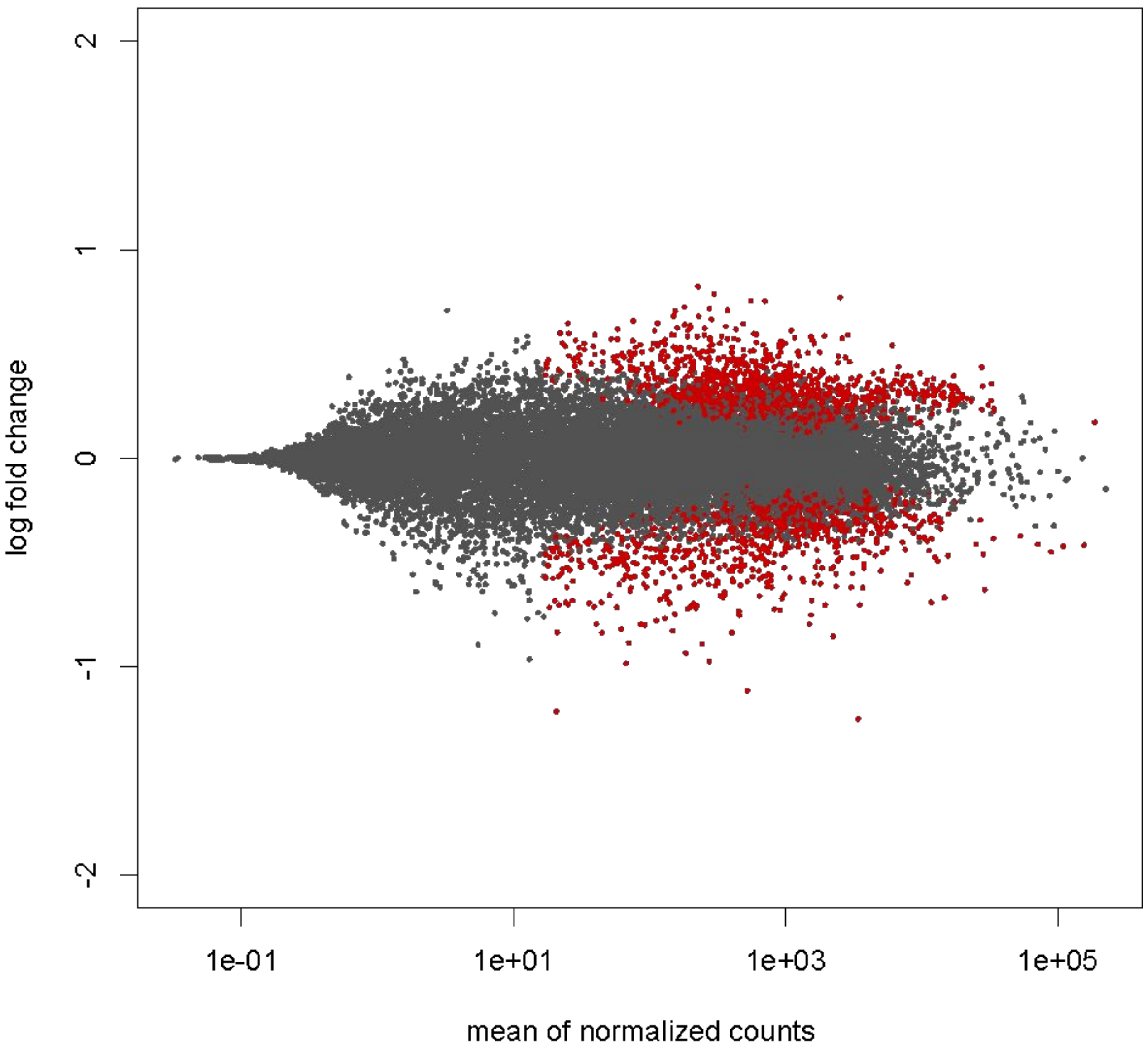


Fig S9. Data smoothing. MA plot showing the mean of normalized counts for all the transcripts. The red points are transcripts with large fold change due to low counts.

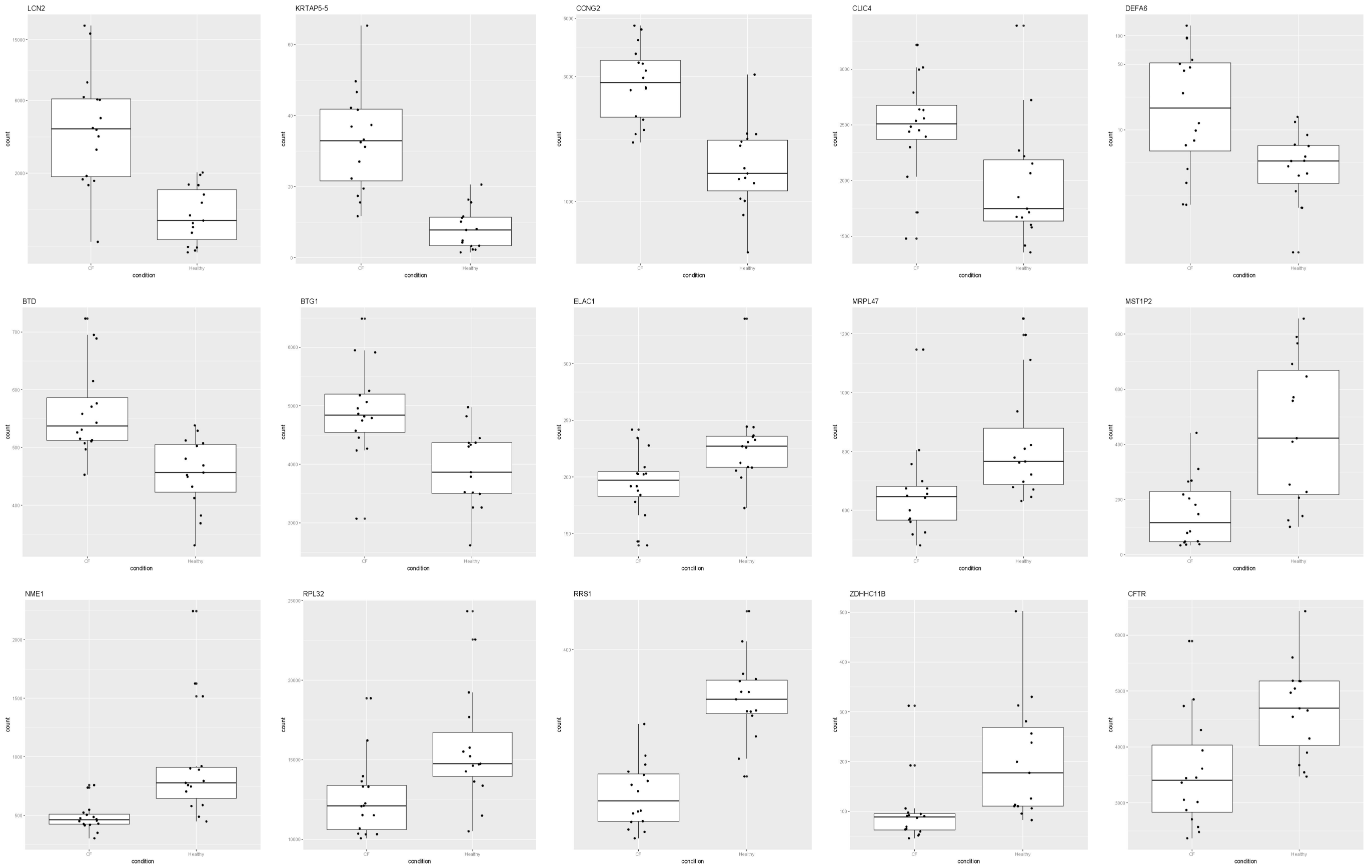


Fig S10. Validation of differentially expressed genes. Box plots of randomly selected DEGs, to show variability of counts between CF and healthy samples.

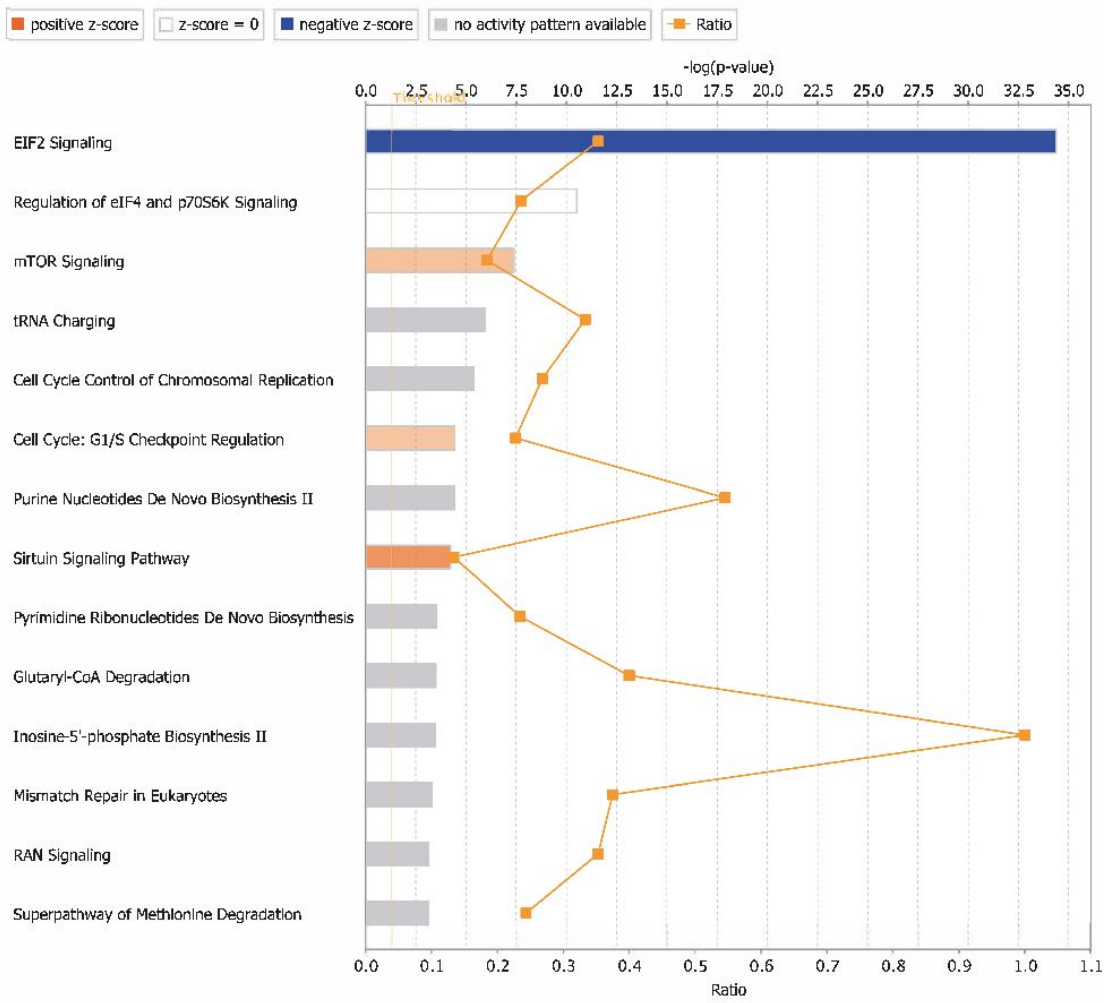


Fig S11. Canonical pathways. Bar plots showing select top pathways based on the negative log p-values for which the DEGs are enriched for. The blue bars are for pathways with negative z-score, while orange bars are for positive z-scores. Furthermore, if the z-score is either zero or has no activity pattern available then those bars are represented by white and grey bars respectively. The orange squares connected by the lines are to show the ratio.

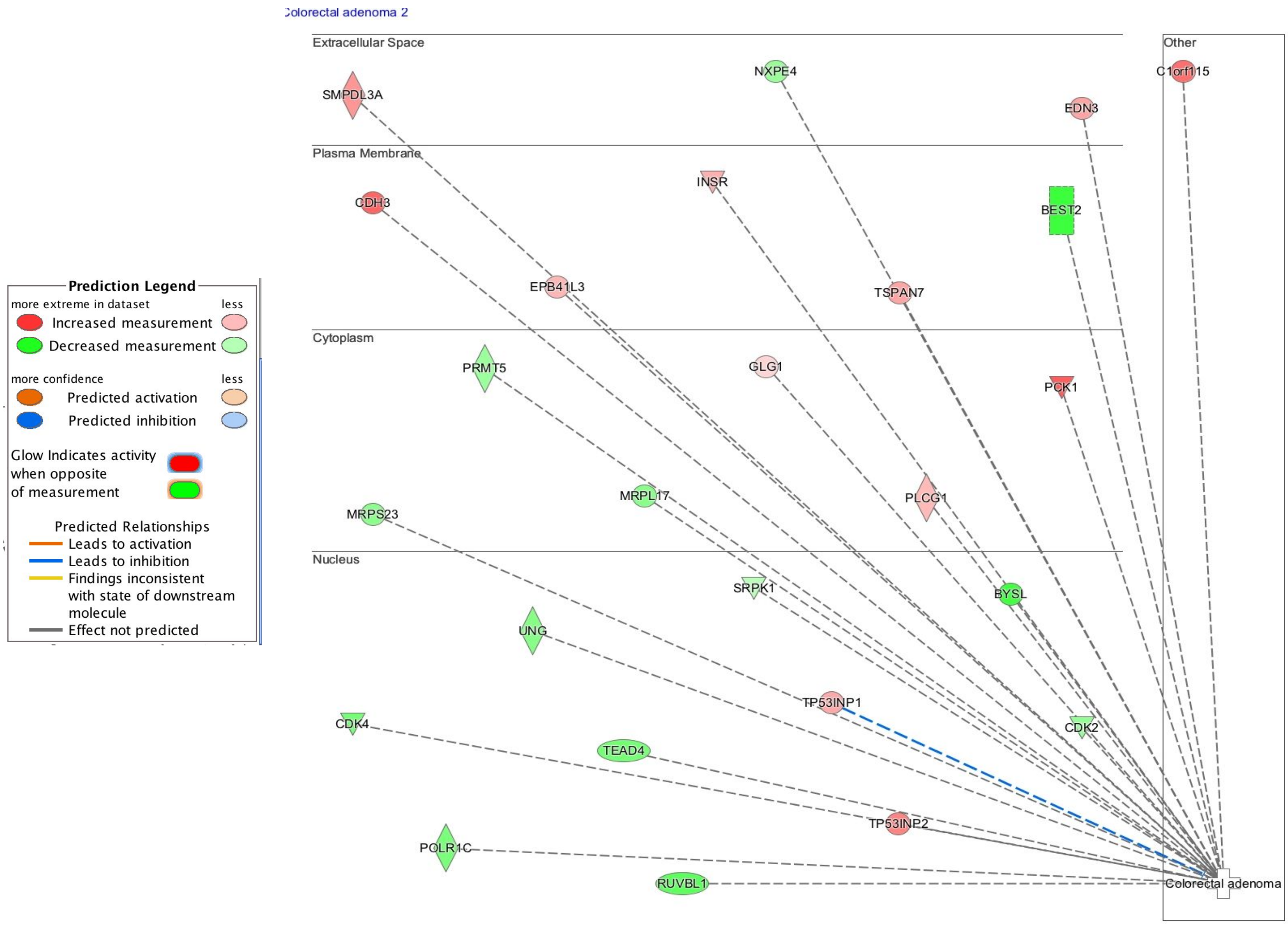


Fig S12. Colorectal gene network. A network of various genes involved in the colorectal cancer disease pathway. The genes have been separated into various bins based on the part of the cell in which they are most active. The upregulated genes in CF are red, while the downregulated genes are green.

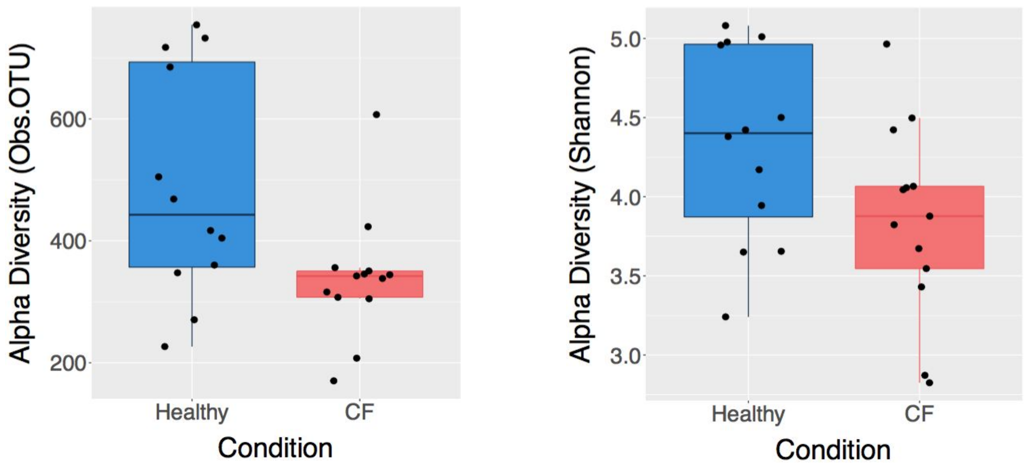


Fig S13: Alpha diversity for observed OTUs and Shannon metrics in CF samples compared to healthy samples (p-value 0.024 and 0.087, respectively, Wilcoxon rank sum test)

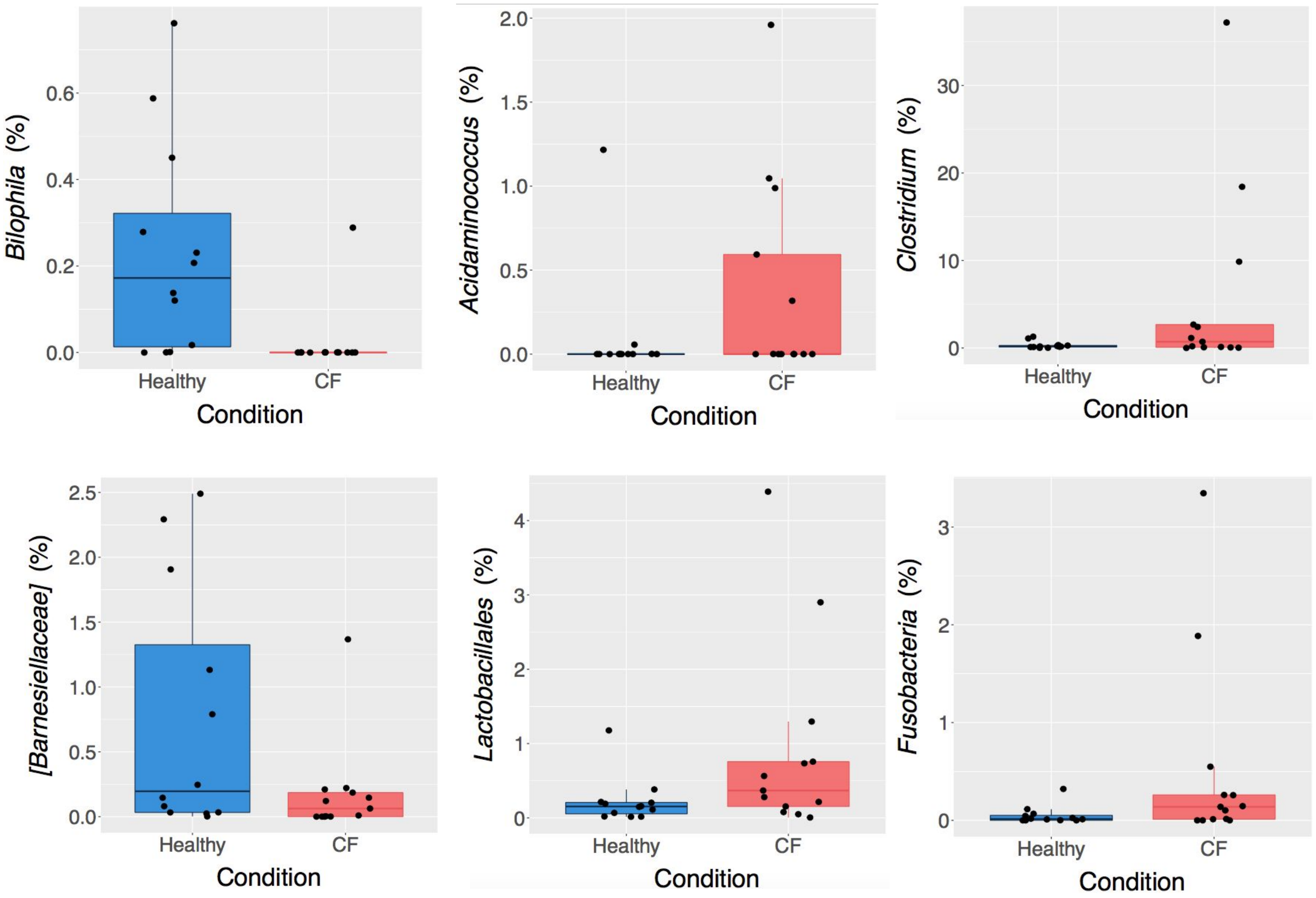


Fig S14: Randomly selected differentially abundant taxa between CF and Healthy conditions (FDR < 0.1, Wald's test). Y-axis denotes percent relative abundance.

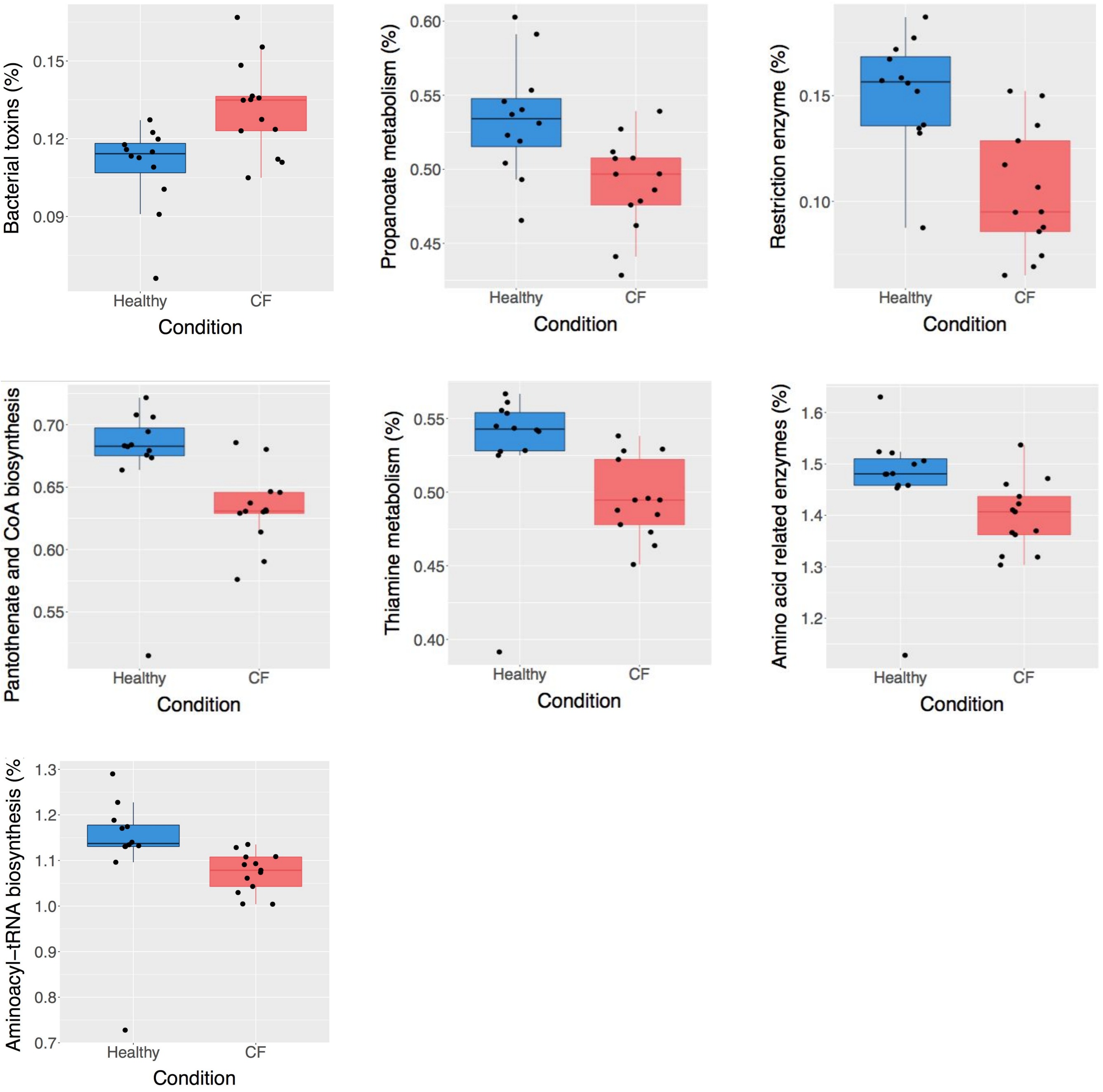


Fig S15: Differentially abundant predicted metabolic pathways in CF samples compared to healthy (FDR < 0.2, Wilcoxon rank sum test). These predicted functional profiles were generated using PICRUSt, where pathways and enzymes were assigned using KEGG database.
